# IP6K1 upregulates the formation of processing bodies by promoting proteome remodeling on the mRNA cap

**DOI:** 10.1101/2020.07.13.199828

**Authors:** Akruti Shah, Rashna Bhandari

**Affiliations:** Laboratory of Cell Signalling, Centre for DNA Fingerprinting and Diagnostics (CDFD), Inner Ring Road, Uppal, Hyderabad 500039, India; Graduate studies, Manipal Academy of Higher Education, Manipal 576104, India

**Keywords:** mRNA decay, mRNA metabolism, P-bodies, translation suppression

## Abstract

Inositol hexakisphosphate kinases (IP6Ks) are ubiquitously expressed small molecule kinases that catalyze the conversion of the inositol phosphate IP6 to 5-IP7. IP6Ks have been reported to influence cellular functions by protein-protein interactions independent of their enzymatic activity. Here, we show that IP6K1 regulates the formation of processing bodies (P-bodies), which are cytoplasmic ribonucleoprotein granules that serve as sites for storage of translationally repressed mRNA. Cells with reduced levels of IP6K1 display a dramatic reduction in the number of P-bodies, which can be restored by the expression of active or catalytically inactive IP6K1. IP6K1 does not localize to P-bodies, but instead facilitates the formation of P-bodies by promoting translation suppression. We demonstrate that IP6K1 is present on ribosomes, where it interacts with proteins that constitute the mRNA decapping complex – the scaffold protein EDC4, activator proteins DCP1A/B, and the decapping enzyme DCP2. IP6K1 also interacts with components of the eIF4F translation initiation complex – the scaffolding protein eIF4G1, the RNA helicase eIF4A2, and the cap binding protein eIF4E. The RNA helicase DDX6 and the eIF4E binding protein 4E-T are known to promote translation suppression to facilitate P-body formation. We show that IP6K1 binds to DDX6 and promotes the interaction of DDX6 and 4E-T with the cap binding protein eIF4E, and also enhances the binding between DDX6 and EDC4, thus acting to suppress mRNA translation and promote mRNA decapping. Our findings unveil IP6K1 as a novel facilitator of proteome remodelling on the mRNA cap, tipping the balance in favour of translation repression over initiation, and thus leading to the formation of P-bodies.

## Introduction

Inositol hexakisphosphate kinases (IP6Ks) catalyse the synthesis of the small signalling molecule 5-diphosphoinositol pentakisphosphate (5PP-IP5 or 5-IP7) from inositol hexakisphosphate (IP6) (1,2). Mammals have three isoforms of IP6Ks – IP6K1, IP6K2, and IP6K3, which exhibit similarities in their C-terminal domains and divergence at their N-termini (2–4). IP6Ks have been shown to influence a variety of cellular and physiological processes in mammals, by virtue of their ability to synthesize 5-IP7, and also via protein-protein interactions that are independent of their catalytic activity (1). The loss of IP6K1 in mice results in male-specific infertility, primarily due to incomplete sperm maturation (5). We have shown that IP6K1 is expressed at high levels in round spermatids, where it is essential for the formation of perinuclear ribonucleoprotein (RNP) granules called chromatoid bodies (6). The absence of chromatoid bodies in *Ip6k1* knockout mice leads to premature translation of key spermiogenic proteins, resulting in defects in spermiogenesis, and concomitant infertility (6). The functional analogue of chromatoid bodies in somatic cells are called processing bodies or P-bodies (7). These cytoplasmic RNP granules are thought to primarily be sites for mRNA storage (8,9), and harbour proteins involved in suppression of mRNA translation (10–12). P-bodies are also enriched in proteins that participate in 5’-3’ mRNA degradation, including proteins responsible for mRNA deadenylation, mRNA decapping, and 5’-3’ exoribonuclease activity (13–17).

Proteins involved in suppression of mRNA translation are essential for the *de novo* formation of P-bodies (18). mRNA translation and mRNA decay are intimately linked - both processes require the mutually exclusive binding of multiprotein complexes to the m^7^GTP mRNA cap. The translation initiation factor eIF4E, and the mRNA decapping enzyme DCP2, compete to bind the mRNA cap (19–21). A proteome exchange at the mRNA cap, where the decapping complex replaces the translation initiation complex, is pre-requisite for P-body formation. Translation suppressors, including the RNA helicase DDX6 and the eIF4E binding protein 4E-T, mediate multivalent interactions that facilitate this proteome switch (10,22–24). However, factors that assist or regulate DDX6/4E-T-mediated mRNP remodelling during the transition from translation to suppression, remain unexplored.

Here, we introduce IP6K1 as a novel factor that promotes P-body formation, independent of its ability to synthesize 5-IP7. Lowering cellular levels of IP6K1 causes a drastic reduction in the abundance of P-bodies. IP6K1 is unique amongst several proteins known to facilitate P-body formation, as it regulates events leading to P-body assembly, but does not localize to P-bodies. We uncover multiple IP6K1 interactions that facilitate the DDX6/4E-T-mediated block in translation and downstream mRNA decapping. We show that IP6K1 localises to ribosomes and facilitates proteome exchange on the mRNA cap as the transcript transitions from active translation to repression, thus promoting the formation of P-bodies.

## Results and Discussion

### Depletion of IP6K1 disrupts P-body formation

We have previously illustrated that IP6K1 is indispensable for the assembly of chromatoid bodies in round spermatids (6). As chromatoid bodies are similar to P-bodies in terms of their function and composition (7), we investigated whether IP6K1 depletion also affects the formation of P-bodies. Bronchiolar epithelial sections from either *Ip6k1*^+/+^ or *Ip6k1*^−/−^ mice were subjected to immunostaining with the P-body marker protein, DCP1A. In sections from *Ip6k1*^+/+^ mice, we observed faint cytoplasmic staining and intense granular staining of DCP1A, indicative of its enrichment in P-bodies (Fig. 1A). However, in *Ip6k1*^−/−^ sections, DCP1A staining was predominantly cytoplasmic, and P-body granules were nearly absent, revealing that IP6K1 is indeed essential for the presence of P-bodies in somatic cells. Next, we examined the human tumour-derived cell lines HeLa and U-2 OS, which normally contain an average of ten P-bodies per cell, and have been used as model systems for the study of P-body assembly and function (8,25). We examined HeLa sh*IP6K1* and U-2 OS sh*IP6K1* cell lines, that harbour shRNA directed against *IP6K1* and display a 70-80% knockdown in IP6K1 expression levels compared with non-targeted control cell lines HeLa NT and U-2 OS NT, respectively ((26), Fig. S1A). DCP1A immunostaining revealed a drastic reduction in the number of P-bodies in IP6K1 depleted cells compared to control cells (Figs. 1B-E). Several proteins that regulate P-body assembly are known to also localize to P-bodies. Since we have earlier reported that IP6K1 is enriched in chromatoid bodies in round spermatids (6), we co-stained U-2 OS cells with antibodies against IP6K1 and DCP1A, to determine whether IP6K1 also localizes to P-bodies. Although IP6K1 displayed some granular cytoplasmic staining, we did not observe any co-localization between IP6K1 and DCP1A in P-bodies (Figs. 1F and G). Even when over-expressed in U2-OS cells, IP6K1 showed diffuse staining throughout the cell, but was not enriched in P-bodies (Fig. S1B). These data indicate that, unlike most proteins involved in P-body assembly, IP6K1 does not localize to P-bodies.

**Figure 1.**
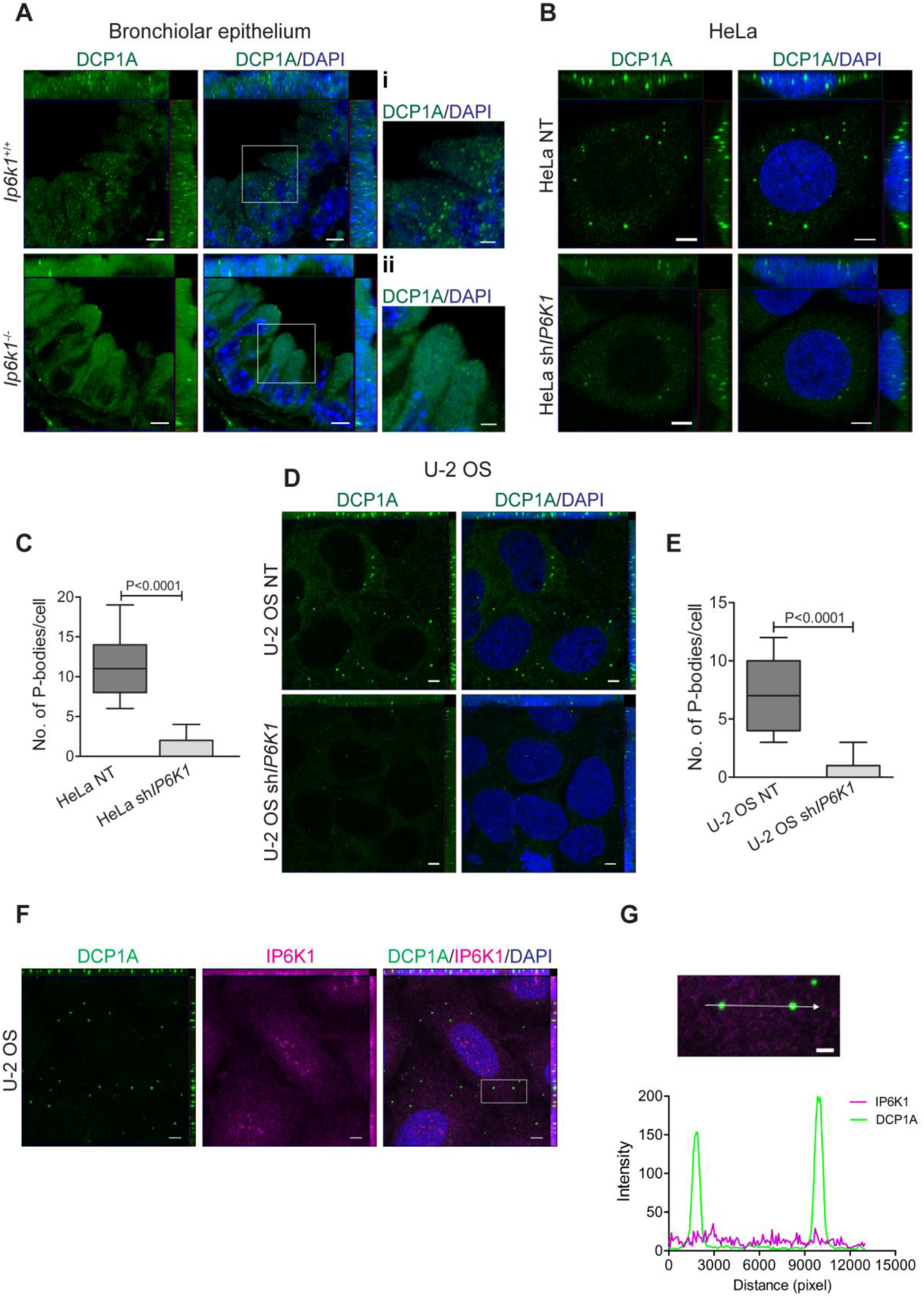
Depletion of IP6K1 reduces the abundance of P-bodies. **(A)** Representative bronchiolar epithelial cross sections from *Ip6k1*^+/+^ and *Ip6k1*^−/−^ mice were stained for P-body marker protein DCP1A (green). Nuclei were stained with DAPI (blue). Boxed areas in (A) were magnified, and are shown as insets (i) and (ii). Inset (i) shows granular P-body staining in *Ip6k1*^+/+^ bronchiolar epithelial cells. Inset (ii) shows diffuse cytoplasmic staining in *Ip6k1*^−/−^ bronchiolar epithelial cells. Scale bar in (A) is 5 μm and in insets (i) and (ii) is 2 μm, N=2. **(B)** Asynchronous HeLa NT and sh*IP6K1* cells were stained with anti-DCP1A antibody (green). Nuclei were stained with DAPI (blue). Scale bar, 5 μm. **(C)** Quantification of the number of P-bodies/cell in (B). Data (mean ± SEM, n = 49 and 36 cells respectively, for HeLa NT and sh*IP6K1* cells) are representative of two independent experiments. **(D)** Asynchronous U-2 OS NT and sh*IP6K1* cells were stained with anti-DCP1A antibody (green). Nuclei were stained with DAPI (blue). Scale bar, 5 μm. **(E)** Quantification of the number of P-bodies/cell in (D). Data (mean ± SEM, n = 109 and 113 cells respectively, for U-2 OS NT and sh*IP6K1* cells) are representative of three independent experiments. **(F)** Localization of IP6K1 (magenta) with DCP1A (green) in U-2 OS cells. Nuclei were stained with DAPI (blue). Scale bar, 5 μm, N=2. **(G)** The boxed region in (F) was magnified and fluorescence intensity profiles were measured along the line drawn. Scale bar, 2 μm. The green and magenta traces denote DCP1A and IP6K1 fluorescence intensities, respectively, and the two green high intensity peaks correspond to P-bodies. Images in (A), (B), (D) and (F) were subjected to uniform ‘levels’ adjustment in the ZEN software to improve visualization. *P* values in (C) and (E) are from a two-tailed unpaired Student’s *t*-test; P ≤ 0.05 was considered significant.

### IP6K1 controls P-body formation independent of its catalytic activity

To determine whether the ability of IP6K1 to synthesize 5-IP7 is essential for the formation of P-bodies, we introduced V5 epitope tagged active IP6K1 (IP6K1-V5) or kinase-dead IP6K1 (IP6K1-V5 K226A) in U-2 OS sh*IP6K1* cells. Expression of either version of IP6K1 restored the ability of U-2 OS sh*IP6K1* cells to form P-bodies, indicating that IP6K1 is required for the presence of P-bodies independent of its catalytic activity (Figs. 2A and B). However, the number of P-bodies per cell was higher in U-2 OS sh*IP6K1* cells expressing active IP6K1 compared with cells expressing the kinase-dead mutant version of IP6K1 (Fig. 2B). To investigate this further, we overexpressed either IP6K1-V5 or IP6K1-V5 K226A in native U-2 OS cells. Overexpression of both these versions of IP6K1 caused a significant increase in the number of P-bodies (Figs. 2C and D). However, as noted in the case of U-2 OS sh*IP6K1* cells, the number of P-bodies in native U-2 OS cells overexpressing active IP6K1 was greater than the number of P-bodies seen in cells overexpressing the inactive enzyme (Fig. 2D). We conclude that IP6K1 protein, irrespective of its enzymatic activity, is sufficient for the formation of P-bodies, but that 5-IP7 acts to increase the number of P-bodies per cell.

**Figure 2.**
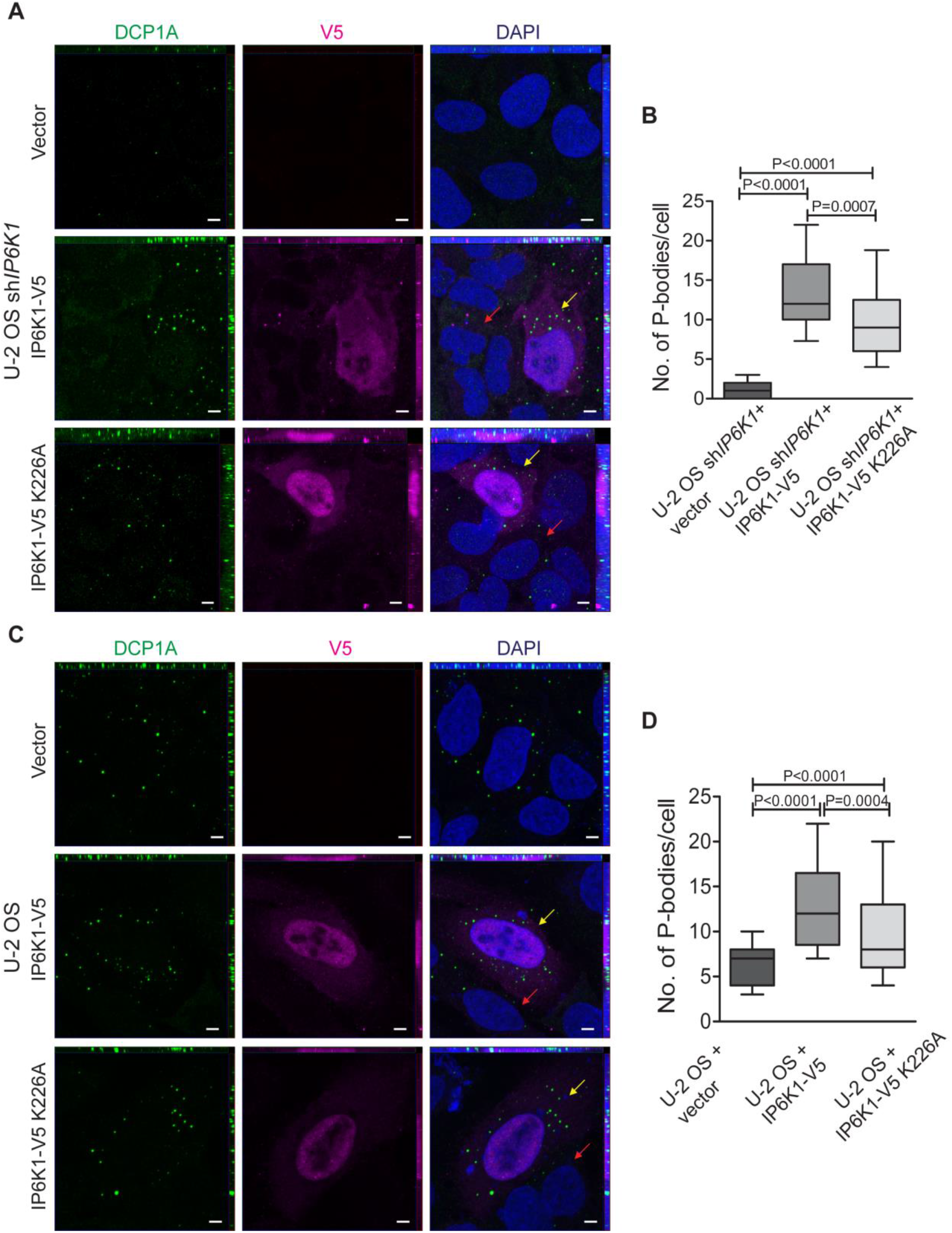
P-body formation does not require IP6K1 catalytic activity. **(A)** Asynchronous U-2 OS sh*IP6K1* cells overexpressing either IP6K1-V5, IP6K1-V5 K226A, or vector control (magenta) were stained with anti-DCP1A antibody (green). Nuclei were stained with DAPI (blue). Transfected and untransfected cells are indicated with yellow and red arrows, respectively. Scale bar, 5 μm. **(B)** Quantification of the number of P-bodies/cell in (A). Data (mean ± SEM, n = 67, 59, and 62 cells respectively, for vector, IP6K1-V5, and IP6K1-V5 K226A overexpressing cells) are from three independent experiments. **(C)** Asynchronous U-2 OS cells overexpressing either IP6K1-V5, IP6K1-V5 K226A, or vector control (magenta) were stained with anti-DCP1A antibody (green). Nuclei were stained with DAPI (blue). Transfected and untransfected cells are indicated with yellow and red arrows, respectively. Scale bar, 5 μm. **(D)** Quantification of the number of P-bodies/cell in (C). Data (mean ± SEM, n = 72, 89, and 66 cells respectively, for vector, IP6K1-V5, and IP6K1-V5 K226A overexpressing cells) are from two independent experiments. Images in (A) and (C) were subjected to uniform ‘levels’ adjustment in the ZEN software to improve visualization. *P* values are from a two-tailed unpaired Student’s *t*-test (B and D); *P* ≤ 0.05 was considered significant.

### IP6K1 depletion causes down-regulation of key mRNA decapping proteins

Since IP6K1 does not localise to P-bodies, it may influence P-body formation via indirect regulation of other key P-body components. We therefore investigated whether lowering the expression of IP6K1 affects the level of key P-body resident proteins. We performed immunoblotting analysis to examine the total cellular levels of components of the mRNA decapping complex, including the scaffold protein EDC4, the decapping enzyme DCP2, and the decapping activator proteins DCP1A and DCP1B. Interestingly, knocking down IP6K1 expression by 70% in U-2 OS sh*IP6K1* cells resulted in a 40-80% reduction in the levels of EDC4, DCP1A/B, and DCP2, when compared to U-2 OS NT cells (Figs. 3A and B). However, this reduction in the level of decapping proteins is independent of any change in the level of their mRNA transcripts (Fig. 3C). We also examined the expression level of other P-body marker proteins including the RNA helicase DDX6, which suppresses mRNA translation, PAN3, which deadenylates mRNA prior to decapping, the 5’-3’ exoribonuclease XRN1, which degrades decapped mRNA, and GW182, a scaffolding protein involved in micro-RNA mediated mRNA degradation. Unlike the decapping protein complex, knocking down IP6K1 expression did not lower the cellular levels of these four proteins (Figs. 3D and E). These findings indicate that IP6K1 is required to sustain the levels of proteins involved in mRNA decapping, but not proteins that function upstream or downstream to decapping. It is possible that the observed reduction in the number of DCP1A stained P-bodies in sh*IP6K1* expressing cells (Figs. 1 B-E) is only a reflection of lower protein levels of DCP1A, but not an actual depletion of cellular P-bodies. To resolve this ambiguity, we performed immunofluorescence analysis on U-2 OS sh*IP6K1* cells using antibodies against XRN1 and DDX6, whose levels are not affected by IP6K1 depletion. U-2 OS sh*IP6K1* cells were found to be consistently devoid of P-bodies even when stained with DDX6 or XRN1 (Figs. S2 A-D)

**Figure 3.**
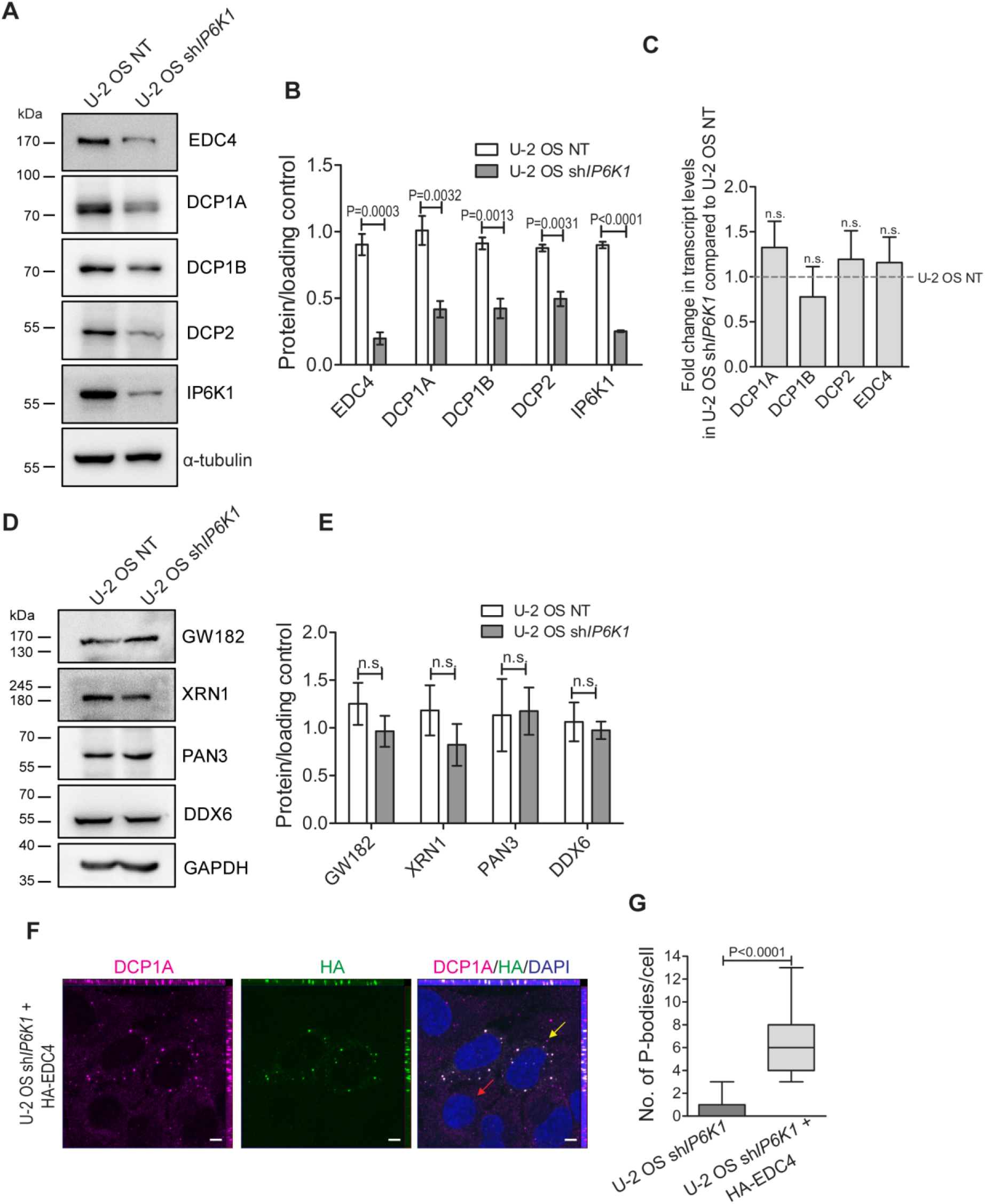
Reduced IP6K1 levels leads to downregulation of mRNA decapping proteins. **(A)** Representative immunoblots showing steady state levels of mRNA decapping proteins in U-2 OS NT and sh*IP6K1* cells. Tubulin was used as a loading control. **(B)** Protein levels from (A) were normalized to their respective loading control. Data are mean ± SEM, N=3 for DCP2, and N=4 for EDC4, DCP1A, DCP1B and IP6K1. **(C)** RT-qPCR analysis of transcripts encoding mRNA decapping proteins that were observed to be downregulated in (A). Values indicate the fold change in transcript levels in U-2 OS sh*IP6K1* cells normalized to U-2 OS NT cells. Data are mean ± SEM, N=3 for DCP1B, and N=4 for EDC4, DCP1A, and DCP2. **(D)** Representative immunoblots showing steady state levels of proteins that function up - or down-stream to mRNA decapping in U-2 OS NT and sh*IP6K1* cells. GAPDH was used as a loading control. **(E)** Protein levels from (D) were normalized to their respective loading control. Data are mean ± SEM, N=3. **(F)** Asynchronous U-2 OS sh*IP6K1* cells overexpressing HA epitope-tagged EDC4 (magenta) were stained with anti-DCP1A antibody (green). Nuclei were stained with DAPI (blue). Transfected and untransfected cells are indicated with yellow and red arrows, respectively. Scale bar, 5 μm. **(G)** Quantification of the number of P-bodies/cell in (F). Data (mean ± SEM n = 56 and 59 cells, respectively, for untransfected and HA-EDC4-transfected cells) are from two independent experiments. Images in (F) were subjected to uniform ‘levels’ adjustment in the ZEN software to improve visualization. *P* values are from a two-tailed unpaired Student’s *t*-test (B, E and G), or one sample *t*-test (C); *P* ≤ 0.05 was considered significant; n.s., not significant *P* > 0.05.

It has been shown that EDC4 provides a scaffold over which DCP1A and DCP2 interact (27). EDC4 downregulation leads to a decrease in DCP2 levels and a reduction in the number of P-bodies (27–29), but a decrease in DCP1A or DCP2 does not affect EDC4 (28,30,31). Since knocking down expression of IP6K1 leads to a reduction in the levels of all three proteins - EDC4, DCP1A, and DCP2, IP6K1 may act primarily via regulating the level of EDC4. To investigate this possibility, we attempted to rescue the depletion of P-bodies in U-2 OS sh*IP6K1* cells by overexpressing EDC4. Immunostaining to detect DCP1A revealed that EDC4 overexpression restores P-body formation, when compared with untransfected cells in the same sample (Figs. 3F and G). Therefore, it is possible that IP6K1 controls P-body assembly by regulating the level of EDC4. If this were the mechanism for upregulation of P-body assembly by IP6K1, the increased abundance of P-bodies observed upon overexpression of IP6K1 (Figs. 2C and D) should be accompanied by an increased expression of mRNA decapping proteins. However, the levels of EDC4, DCP1A and DCP2 remained unaltered upon overexpression of either active or inactive IP6K1 (Figs. S2E and F). This data indicates that IP6K1 does not regulate P-body formation merely by controlling the expression level of mRNA decapping proteins, and points to an additional layer of regulation of P-body formation by IP6K1.

### IP6K1 interacts with the mRNA decapping complex on ribosomes

To further explore the mechanism by which IP6K1 controls P-body dynamics, we first sought to check if IP6K1 binds any of the mRNA decapping proteins. IP6K1 was able to specifically pull down endogenous DCP1A, DCP2 and EDC4 (Fig. 4A). These interactions persisted even after RNAse A treatment of cell lysates, indicating that the binding of IP6K1 to the mRNA decapping complex is RNA-independent (Fig. S3A), and most likely a direct protein-protein interaction. The RNA helicase DDX6, a component of the mRNA decapping complex, is known to promote P-body assembly by repressing translation initiation (18). We noted that IP6K1 can also interact with endogenous DDX6 (Fig. 4B).

**Figure 4.**
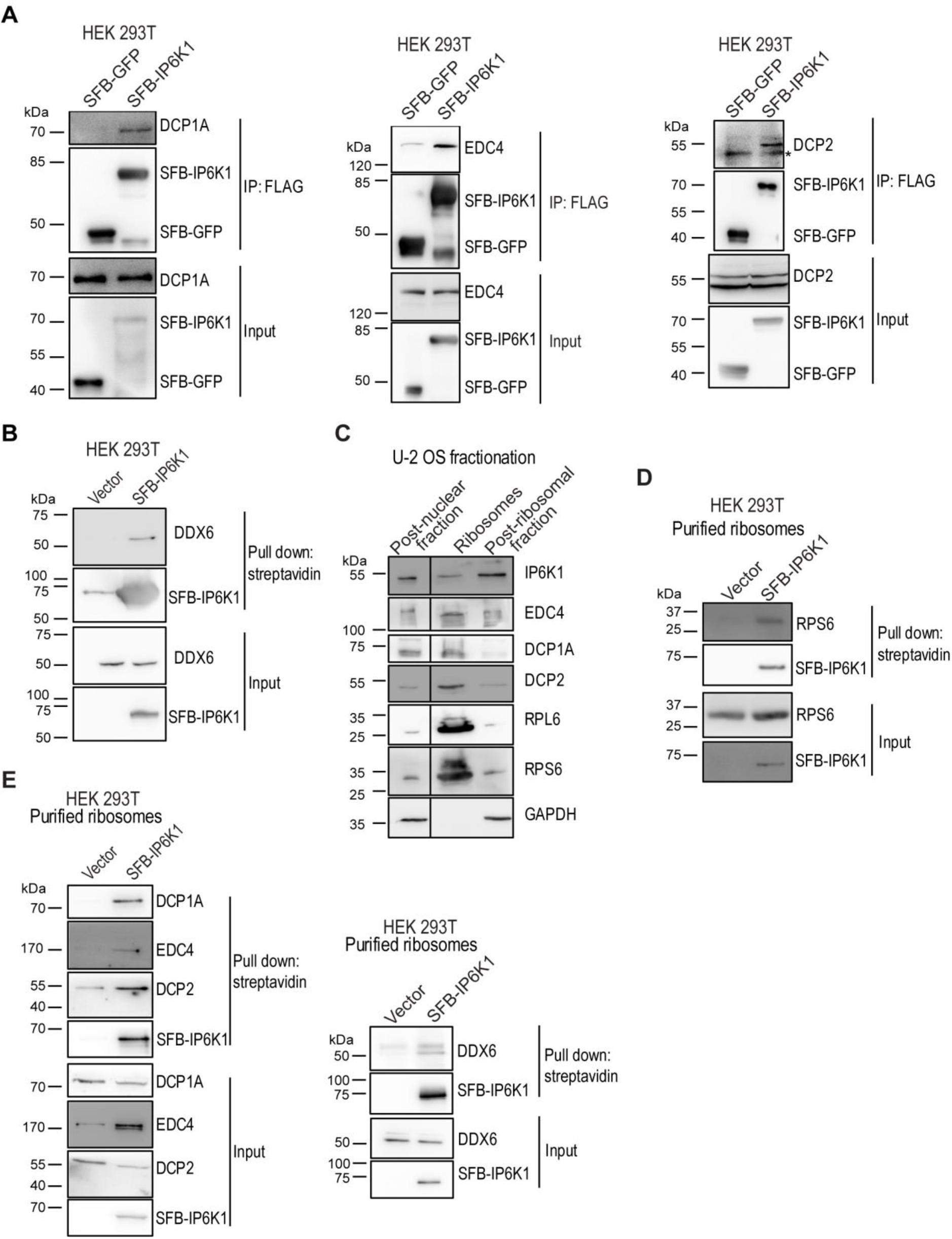
IP6K1 interacts with the mRNA decapping complex on ribosomes. **(A)** Representative immunoblots showing co-immunoprecipitation of endogenous DCP1A, EDC4, and DCP2 with SFB-tagged IP6K1. SFB-IP6K1 or SFB-GFP were transiently overexpressed in HEK293T cells, immunoprecipitated with an anti-FLAG antibody, and probed to detect DCP1A, EDC4, or DCP2. The SFB tag was detected using an anti-FLAG antibody (N=3). An asterisk (*) indicates bands corresponding to heavy chain of the antibody used for immunoprecipitation. (**B**) Representative immunoblots showing co-immunoprecipitation of endogenous DDX6 with SFB-tagged IP6K1. SFB-IP6K1 was transiently over-expressed in HEK293T cells, pulled-down with streptavidin sepharose beads, and probed to detect DDX6. The SFB tag was detected using an anti-FLAG antibody (N=3). **(C)** Representative immunoblots of subcellular fractions of U-2 OS cells prepared by ultracentrifugation, to detect endogenous IP6K1, EDC4, DCP1A, and DCP2. Enrichment of ribosomes was marked by the presence of RPL6 and RPS6, and GAPDH was detected to rule out cytoplasmic contamination in the purified ribosomes (N=4). Vertical line indicates removal of non-essential lanes from a single original gel to improve visualisation. **(D)** Representative immunoblots showing co-immunoprecipitation of RPS6 with SFB-IP6K1. Purified ribosomes isolated from HEK293T cells transiently overexpressing SFB-IP6K1 were subjected to pull-down with streptavidin sepharose beads, and probed to detect RPS6 (N=2). **(E)** Representative immunoblots showing co-immunoprecipitation of decapping proteins with SFB-IP6K1 in purified ribosomes. The ribosomal fraction isolated from HEK293T cells transiently overexpressing SFB-IP6K1 was subjected to pull-down with streptavidin beads, and probed to detect DCP1A, EDC4, DCP2 and DDX6 (N=3).

Our data show that IP6K1 specifically interacts with P-body-enriched mRNA decapping proteins, but that IP6K1 is itself not localized to P-bodies. Therefore, this protein-protein interaction is likely to occur at another sub-cellular compartment. In budding yeast and plants, several proteins involved in mRNA decapping and degradation are shown to localize to ribosomes on the endoplasmic reticulum membrane, and facilitate co-translational degradation of stalled mRNAs (32–34). To examine the possibility that IP6K1 interacts with the mRNA decapping machinery on ribosomes, we first checked if decapping proteins and IP6K1 localize to ribosomes in mammalian cells. We fractionated U-2 OS and 293T cell extracts, and obtained a pure ribosomal fraction, as evidenced by the absence of the cytoplasmic marker GAPDH, and enrichment of small and large ribosomal subunit proteins RPS6 and RPL6 respectively (Fig. 4C and S3B). Interestingly, we detected the presence of both IP6K1, and the mRNA decapping complex (DCP1A, DCP2, and EDC4) in purified ribosomes obtained from U-2 OS and 293T cells (Fig. 4C and S3B). To verify that the interaction between IP6K1 and mRNA decapping proteins occurs on ribosomes, we purified the ribosomal fraction from HEK293T cells expressing tagged IP6K1. Co-precipitation of the ribosomal marker RPS6 with IP6K1 confirmed that IP6K1 is indeed present in purified ribosomes (Fig. 4D). The endogenous decapping proteins DCP1A, DCP2, EDC4 and DDX6 were also pulled down with IP6K1 from the ribosomal fraction (Fig. 4E). Together, these data reveal for the first time, that the mRNA decapping complex is enriched in mammalian ribosomes, and that IP6K1 interacts with these proteins on ribosomes.

### IP6K1 facilitates the interaction between eIF4E and 4E-T/DDX6 to promote translation inhibition and mRNA decapping

An early event in the formation of P-bodies is the exchange of proteins bound to translationally stalled mRNA – proteins responsible for active translation are replaced with proteins involved in mRNA degradation. We hypothesized that the presence of IP6K1 in ribosomes may facilitate this proteome switch. Since IP6K1 interacts with the mRNA decapping complex, we examined whether it also binds to the translation initiation complex eIF4F. Pull down of SFB-tagged IP6K1 revealed a specific interaction with all three members of the eIF4F translation initiation complex - the cap binding protein eIF4E, the RNA helicase eIF4A2, and the scaffolding protein eIF4G1 (Fig. 5A). In translationally stalled mRNA, eIF4E bound to the m^7^GTP cap is replaced with the decapping enzyme DCP2 (21). To determine if IP6K1 facilitates this exchange, we first examined whether endogenous IP6K1 interacts with the eIF4F complex on the mRNA cap. Incubation of U-2 OS cell extract with immobilised m^7^GTP cap analogue revealed that IP6K1 is bound to the mRNA cap, along with the cap binding protein eIF4E (Fig. 5B). To ensure that this pull-down is specific, we treated the cell lysate with monomethylated cap analogue (m^7^GpppG), to competitively deplete eIF4E bound to m^7^GTP beads. The amount of both eIF4E and IP6K1 bound to m^7^GTP beads was reduced in the presence of m^7^GpppG, indicating that IP6K1 is specifically recruited to the mRNA cap (Fig. 5B and C).

**Figure 5.**
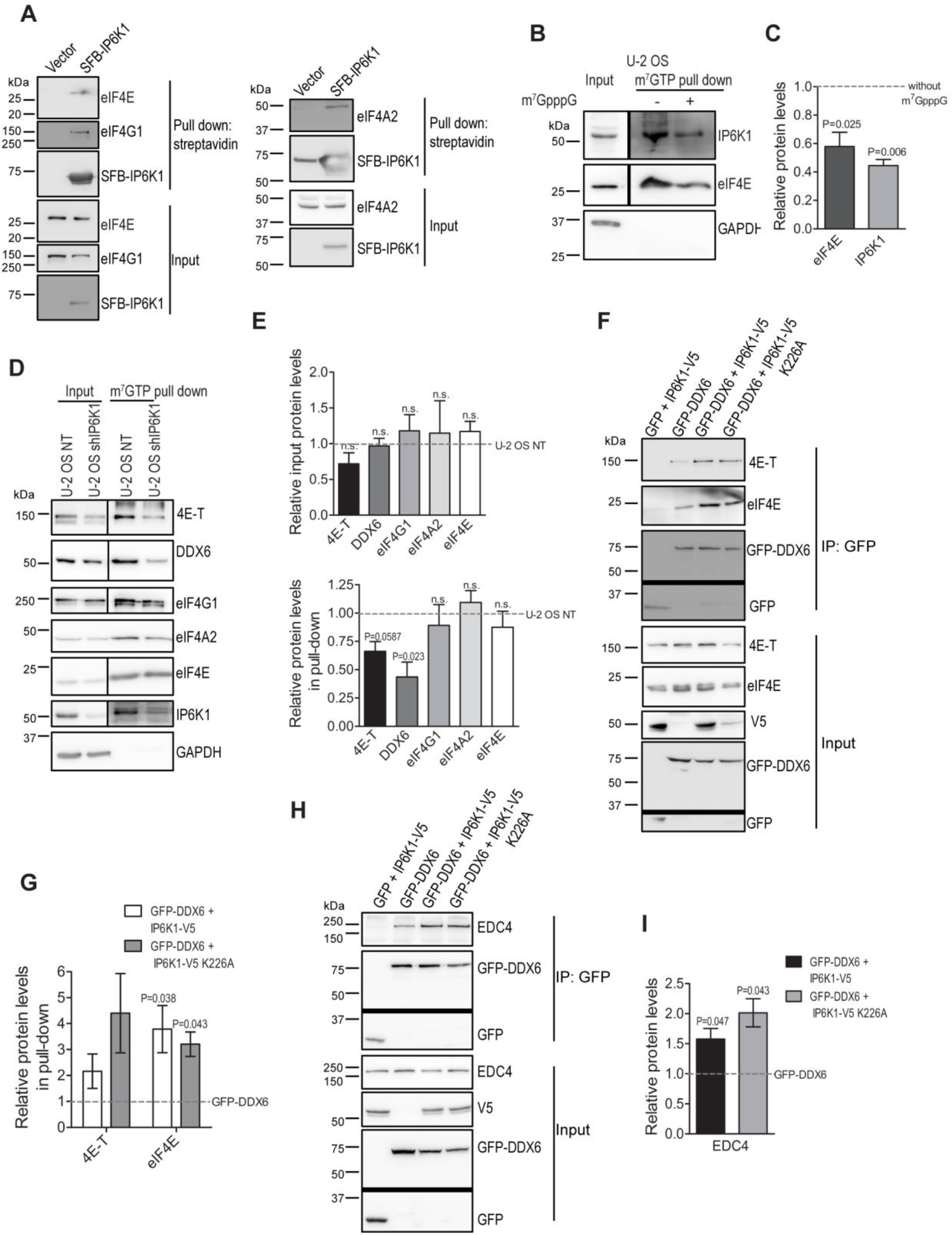
IP6K1 binds the translation initiation complex eIF4F at the mRNA cap. **(A)** Representative immunoblots showing interaction between SFB-tagged IP6K1 and the eIF4F complex. SFB-IP6K1 was transiently overexpressed in HEK293T cells, pulled-down with streptavidin sepharose beads, and probed to detect eIF4E, eIF4G1, or eIF4A2 (N=3). **(B, C)** A U-2 OS cell extract was incubated with m^7^GTP–sepharose beads in the presence or absence of the cap analogue m^7^GpppG, and subjected to immunoblotting to detect IP6K1 and eIF4E. Vertical line indicates separation of input and pull-down lanes from a single blot, that were subjected to differential ‘auto-contrast’ adjustment for better visualization of bands in the pull-down lane. Bar graphs show mean fold change ± SEM in the extent of pull-down of IP6K1 and eIF4E in the presence of the cap analogue when compared to the absence of cap analogue (N=3). **(D, E)** Representative immunoblots of proteins pulled down from U-2 OS NT and sh*IP6K1* cell extracts by m^7^GTP–sepharose beads. Vertical line indicates separation of input and pull-down lanes from a single blot, that were subjected to differential ‘auto-contrast’ adjustment for better visualization of bands in the pull-down lane. Bar graphs show mean fold change ± SEM in the extent of pull-down of the indicated proteins from U-2 OS sh*IP6K1* cells when compared to U-2 OS NT cells (N=3 for 4E-T and eIF4A2, and N=4 for DDX6, eIF4G1 and eIF4E). **(F-I)** Representative immunoblots showing the effect of overexpression of V5 epitope-tagged IP6K1 or IP6K1 K226A on the extent of pull-down of endogenous 4E-T and eIF4E (F, G), or EDC4 (H, I) with GFP-tagged DDX6, from HEK293T cell extracts. Bar graphs show the mean fold change ± SEM in the extent of co-immunoprecipitation when either IP6K1 or IP6K1 K226A was co-overexpressed with DDX6, compared with DDX6 overexpression alone (N=3 for eIF4E and EDC4, and N=2 for 4E-T). Horizontal lines indicate division of a single membrane into two parts that were immunoblotted separately, but were subjected to identical exposure during image acquisition. *P* values are from a one sample *t*-test (C, E, G and I); *P* ≤ 0.05 was considered significant; n.s., not significant *P* > 0.05.

The eIF4E transporter protein 4E-T, supresses translation upon binding to eIF4E, and is indispensable for the formation of P-bodies (10). To investigate if IP6K1 regulates 4E-T binding to eIF4E, we pulled down eIF4E from U-2 OS NT and sh*IP6K1* cells using immobilised m^7^GTP cap analogue. Although the expression of 4E-T protein remains unaltered, we observed a significant reduction in the interaction of 4E-T with eIF4E in cells with lower levels of IP6K1 (Fig. 5D and E). The level of expression of eIF4E, eIF4G1, and eIF4A2, or their binding to the mRNA cap analogue, was not affected by IP6K1 (Fig. 5D and E). 4E-T is known to interact with the RNA helicase DDX6 (11,18), and this interaction is required for suppression of translation prior to the formation of P-bodies. We noted that the m^7^GTP cap analogue pulls down DDX6 from a U-2 OS cell extract, indicating that DDX6 and 4E-T are both bound simultaneously to eIF4E (Fig. 5D). The binding of DDX6 to the m^7^GTP cap analogue was also reduced in cells expressing lower levels of IP6K1 (Fig. 5D and E). These data suggest that IP6K1 promotes the binding of the DDX6/4E-T complex to eIF4E on the mRNA cap. To further examine this possibility, DDX6 fused with GFP was immunoprecipitated in the presence or absence of overexpressed IP6K1 in HEK293T cells. The expression of active or catalytically inactive IP6K1 led to a significant increase in the binding of both eIF4E and 4E-T to DDX6 (Fig. 5F and G). Thus, independent of its catalytic activity, IP6K1 promotes the interaction of translation suppressors essential for P-body formation with the translation initiation factor eIF4E. DDX6 is known to have multivalent interactions with both translation suppressors, and mRNA decapping proteins (18). To determine whether IP6K1 also regulates the binding of the mRNA decapping complex to DDX6, we examined the co-immunoprecipitation of EDC4 with DDX6. There was a notable increase in the level of endogenous EDC4 interacting with GFP-DDX6 upon the overexpression of either active or catalytically inactive IP6K1 (Fig. 5H and I). Together, these data illustrate that IP6K1 facilitates proteome remodelling on the mRNA cap, promoting the transition from translation initiation to suppression, and ultimately to mRNA decapping.

Lastly, we performed functional assays to monitor the effect of IP6K1 on suppression of translation and mRNA decapping. The interaction between DDX6 and 4E-T has been shown to be important for micro RNA (miRNA)-mediated suppression of translation, and the subsequent formation of P-bodies (11). Since IP6K1 promotes the binding of 4E-T to DDX6 (Fig. 5F and G), we investigated whether a decrease in cellular IP6K1 levels affects miRNA-mediated mRNA suppression. We employed a miRNA reporter assay that utilizes a Renilla luciferase coding sequence with a let7a miRNA binding site in the 3’UTR, which is either a site for perfect complementary binding (RL-perf) or binding with three sets of mismatches (pRL-3XB) (Fig. 6A). When bound to let7a miRNA, the RL-perf reporter mRNA is cleaved endonucleolytically, but the pRL-3XB mRNA is translationally stalled (35). Any decrease in let7a miRNA mediated mRNA cleavage or translation suppression, is recorded as an increase in Renilla luciferase activity. We noted no change in endonucleolytic cleavage of the reporter mRNA in HeLa sh*IP6K1* cells compared with HeLa NT cells (Fig. 6A). However, Renilla luciferase expression by the pRL-3XB reporter was significantly higher in HeLa sh*IP6K1* cells compared with HeLa NT cells, indicating that IP6K1 promotes miRNA-mediated translation suppression. Since IP6K1 overexpression leads to increased binding of the mRNA decapping complex scaffold protein EDC4 to DDX6, we examined whether this strengthened interaction leads to an increase in mRNA decapping. We co-overexpressed GFP-tagged decapping enzyme DCP2 with IP6K1 in HEK293T cells. Immunoprecipitated GFP-DCP2 was incubated with radiolabelled m^7^GTP-capped mRNA, the products were resolved by thin layer chromatography, and the magnitude of decapping was monitored by densitometry analysis of the decapping product, m^7^GDP. We observed a 30% increase in the extent of decapping when GFP-DCP2 was co-expressed with IP6K1 as compared to the vector control (Fig. 6B and C). Collectively these data show that IP6K1 acts independently of its catalytic activity, via protein-protein interactions, to influence mRNA metabolism by downregulating mRNA translation and upregulating mRNA decapping.

**Figure 6.**
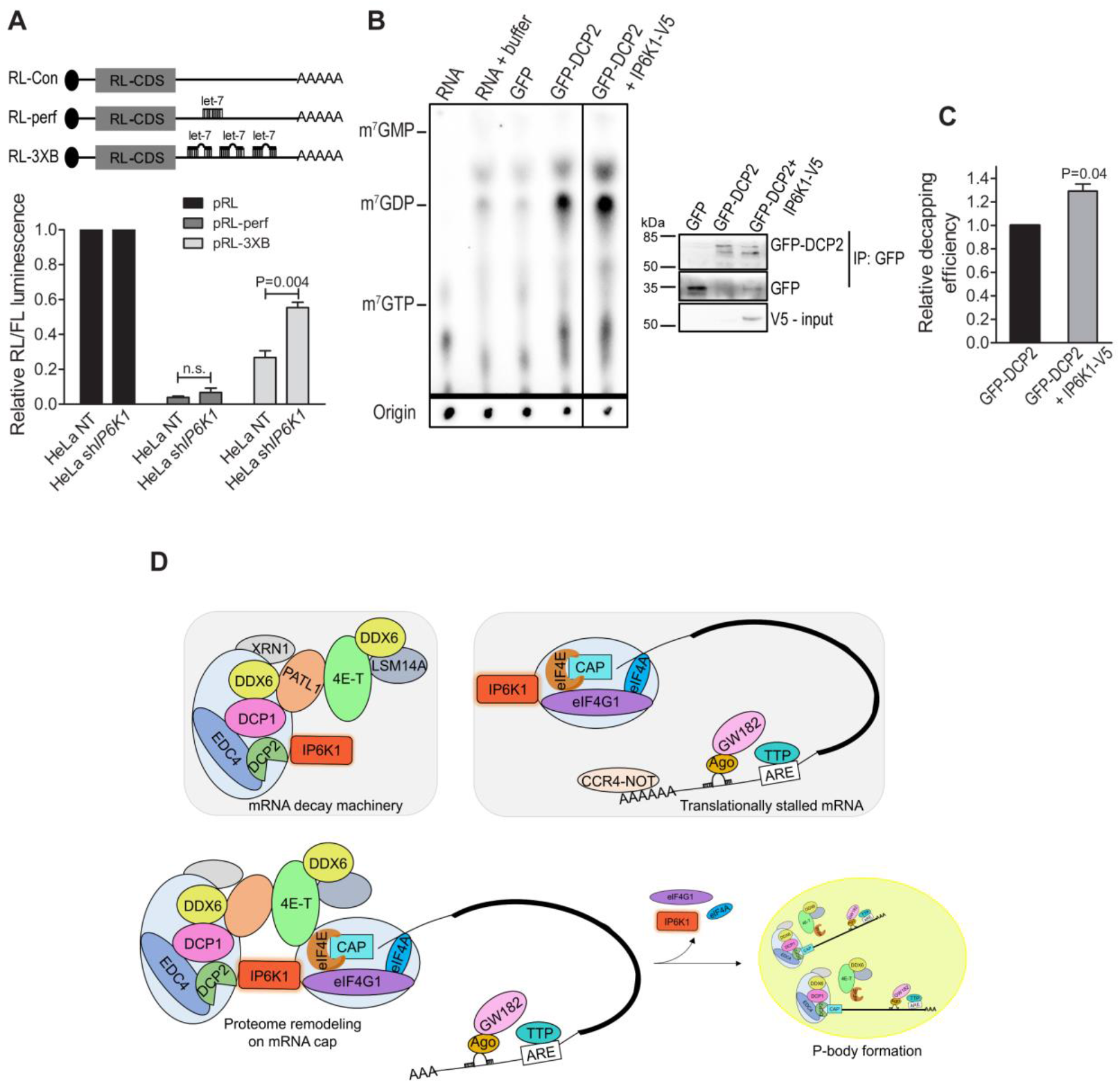
IP6K1 promotes translation suppression and mRNA decapping. **(A)** HeLa NT and sh*IP6K1* cells were transfected with Renilla luciferase (RL) reporter constructs - pRL-control (no let-7a binding site in the 3’UTR), pRL-perf (perfect complimentary let7a binding site in the 3’UTR), or pRL-3xB (3 imperfect let-7a binding sites in the 3’UTR). Firefly luciferase (FL) was co-transfected to serve as an internal control. Relative RL/FL luminescence ratio was calculated, and normalised to the pRL control for each cell line. Data are mean ± SEM, N=3. *P* values are from a two-tailed unpaired Student’s *t*-test; *P* ≤ 0.05 was considered significant; n.s., not significant *P* > 0.05. **(B)** Representative autoradiogram showing the effect of IP6K1 on DCP2-mediated mRNA decapping. Either GFP or GFP-tagged DCP2 were transiently overexpressed in HEK293T cells, with or without V5 epitope tagged-IP6K1. Proteins were immunoprecipitated with anti-GFP antibody and, half of the immunoprecipitate was incubated with ^32^P cap labelled mRNA. The reaction mix was resolved by thin layer chromatography, and the mRNA substrate and hydrolyzed m^7^GDP cap were visualized by autoradiography. Non-radioactive standards were run in parallel and visualized by UV shadowing. The vertical line indicates removal of non-essential lanes from a single original chromatogram for better depiction, and the horizontal line indicates the two parts that were subjected to differential auto-contrast adjustment for better visualization of the m^7^GDP product. The other half of the immunoprecipitate was subjected to immunoblotting with anti-GFP and anti-V5 antibodies. **(C)** Quantification of (B). The extent of decapping by DCP2 in the presence of IP6K1-V5, was quantified and normalized to the extent of decapping by DCP2 alone. Data (mean ± SEM, N=3), were analysed using a one-sample *t*-test. *P* ≤ 0.05 was considered significant. **(D)** IP6K1 facilitates proteome remodelling on the mRNA cap to promote translation stalling. IP6K1 associates with the mRNA decapping complex and the translation initiation complex (these complexes are shown inside light blue capsules to denote that IP6K1 exerts either direct or indirect binding with all of the indicated components). The ability of IP6K1 to interact with both the complexes allows it to mediate multi-protein interactions that result in translation stalling. Specifically, IP6K1 upregulates the association between the translation suppression complex 4-ET/DDX6 and the cap binding protein eIF4E, and binding of the decapping enhancer EDC4 with DDX6.

### IP6K1 promotes proteome remodeling on the mRNA cap to upregulate P-body formation

Our work reveals that IP6K1 is a key player in the regulation of mRNA turnover and P-body formation. Unlike its involvement in the assembly of chromatoid bodies as an integral component, IP6K1 promotes P-body formation without actually localizing to P-bodies. It does this primarily by facilitating translation stalling of mRNAs destined for P-bodies (Fig. 6D). IP6K1 is bound to ribosomes, where it interacts with the mRNA decapping machinery, including EDC4, DCP1A/B, and DCP2, and the RNA helicase DDX6. We also show that IP6K1 binds the translation initiation complex (eIF4G1, eIF4E and eIF4A), and promotes the interaction of DDX6 and 4E-T with the cap binding protein eIF4E. The interaction between DDX6 and 4E-T, and the binding of 4E-T to eIF4E, have been shown to enhance the suppression of translation (11). Our data therefore suggests that IP6K1 facilitates proteome remodeling on the mRNA cap, to tip the scales in favour of translation stalling over translation initiation, and feed P-body formation (Fig. 6D). Our study also shows that a decrease in IP6K1 leads to a reduction in the levels of mRNA decapping proteins EDC4, DCP1A/B and DCP2 (Figs. 3A, B). It is possible that the decrease in translation suppression in cells expressing sh*IP6K1* has a cascading effect on the stability of decapping proteins. Alternatively, IP6K1, or its enzymatic product 5-IP7, may indirectly regulate the levels of decapping proteins by influencing their expression or stability, independent of the effect of IP6K1 on translation suppression. Nonetheless, a reduction in the level of the decapping scaffold protein EDC4, combined with reduced translation suppression, results in a decrease in P-bodies in cells with reduced IP6K1 expression.

Our work shows that the influence of IP6K1 on the formation of P-bodies is largely independent of its catalytic activity. Active or kinase-dead versions of IP6K1 are able to upregulate protein-protein interactions that facilitate translation suppression (Figs. 5D and E). Overexpression of either active or inactive forms of IP6K1 can induce the formation of P-bodies in U-2 OS cells (Fig. 2). However, we do observe a statistically significant difference in the number of P-bodies that accumulate upon over-expression of these two forms of IP6K1, with the catalytically inactive form of IP6K1 inducing fewer P-bodies compared with the active form (Figs. 2B and D). This data correlates with the recent demonstration by Shears and colleagues that an increase in intracellular levels of 5-IP7 upregulates the formation of P-bodies (Stephen Shears, personal communication). The Shears laboratory reports that 5-IP7 competes with the m^7^GTP mRNA cap for binding to NUDT3, a diphosphoinositol polyphosphate phosphohydrolase, which is known to decap a subset of mRNAs independent of DCP2. In cells with increased 5-IP7, there is an accumulation of NUDT3 mRNA substrates, and a consequent elevation in the abundance of P-bodies. IP6K1 has thus emerged as a unique player in the regulation of mRNA turnover and P-body dynamics. It influences the formation of P-bodies via two different mechanisms – by 5-IP7-dependent inhibition of NUDT3-mediated mRNA decapping, and by enzymatic activity independent facilitation of protein-protein interactions that promote DCP2-dependent mRNA decapping.

## Materials and Methods

### Reagents

All chemicals were procured from Sigma-Aldrich, unless specified otherwise. The primary antibodies used in this study for immunofluorescence (IF), immunoblotting (IB) or immunoprecipitation (IP), along with antibody dilution for each application, and supplier (including catalogue number) are as follows: anti-IP6K1 (Sigma-Aldrich, HPA040825; IF 1:500; IB 1:2000), anti-GAPDH (Sigma-Aldrich, G8795; IB 1:10,000), anti-α-tubulin (Sigma-Aldrich, T9026; IB 1:10,000), anti-V5 tag (Thermo Fisher Scientific, R960-25; IB 1:5000, IP 1μg, and Abcam, Ab53418; IF 1:800), anti-GFP (Invitrogen, A11122; IB 1:5000, IP 2μg), anti-HA (Sigma-Aldrich, H6908; IF: 1:500); anti-DCP1A (Abcam, Ab47811; IF 1:400, IB 1:200); anti-DCP1B (Abcam, Ab222313; IB 1:1000); anti-DCP2 (Sigma-Aldrich, D6319; IB 1:2000); anti-EDC4 (Abcam, Ab72408; IB 1:2000); anti-PAN3 (Abcam, Ab189413; IB 1:2000); anti-GW182 (Abcam, Ab156173; IB 1:1000); anti-DDX6 (Novus Biologicals, NB200-191; IF 1:400, IB 1:2000); anti-XRN1 (Abcam, Ab70259; IB 1:2000); anti-eIF4G1 (Cell Signaling Technology, 8701S; IB 1:2000); anti-eIF4E (Novus Biologicals, MAB3228; IB 1:1000); anti-eIF4A2 (Novus Biologicals, NBP2-24529; IB 1:1000). Restriction enzymes were purchased from New England Biolabs. The dual luciferase reporter assay kit was purchased from Promega (E1910). Dulbecco’s Modified Eagle’s Medium (DMEM), Fetal Bovine Serum (FBS) and other cell culture reagents were from Thermo Fisher Scientific. m^7^GTP cap analogue beads (AC-155S), m^7^GTP (NU-1122S), m^7^GDP (NU-1134S), m^7^GMP (NU-1135S), and m^7^GpppG cap analogue (NU-852S) were purchased from Jena Bioscience. NuPAGE 4-12% Bis-Tris gels were from Thermo Fisher Scientific. [α-^32^P]GTP was procured from JONAKI/BRIT. The in vitro transcription and mRNA capping kit was purchased from CellScript (C-MSC11610).

### Plasmids

Human IP6K1 cDNA was sub-cloned into the NheI and BamHI restriction enzyme sites in the pCDNA 3.1 cMYC-V5 expression plasmid (https://www.biorxiv.org/content/10.1101/2020.02.12.945634v1), such that cMYC is replaced by IP6K1, to obtain a C-terminally V5 epitope-tagged IP6K1 plasmid. A catalytically inactive version of human IP6K1 was prepared by using this plasmid to substitute Lys226 with Ala by site-directed mutagenesis and overlap extension PCR, and confirmed by DNA sequencing. Mouse IP6K1 was sub-cloned into SFB (S-protein/FLAG/SBP) - tagged destination vector using the Gateway cloning strategy (Invitrogen). Catalytically inactive IP6K1 (K226A) was generated by site-directed mutagenesis and overlap extension PCR, and cloned into SFB-tagged destination vector. pT7-EGFP-C1-HsRCK (Addgene plasmid # 25033; GenBank ID NM_004397), pCIneo-lambdaN-HA-HsEDC4 (Addgene plasmid # 66597; GenBank ID NM_014329.4) and pT7-EGFP-C1-HsDCP2 (Addgene plasmid # 25031; GenBank ID AY146650) were gifts from Elisa Izaurralde (Max Planck Institute for Developmental Biology, Tübingen, Germany).

### Cell lines and transfection

All cell lines were grown in a humidified incubator with 5% CO_2_ at 37°C in Dulbecco’s modified Eagle’s medium (DMEM) supplemented with 10% fetal bovine serum, 1 mM L-glutamine, 100 U/mL penicillin, and 100 μg/mL streptomycin. HeLa NT and sh*IP6K1* cell lines have been described earlier (36). U-2 OS NT and sh*IP6K1* cell lines were generated similarly. Briefly, lentiviral vectors (pLKO.1) encoding either a non-targeting shRNA (SHC016, Sigma-Aldrich) or shRNA directed against human *IP6K1* (TRC0000013508, Sigma-Aldrich) were co-transfected with VSV-G and psPAX2 (a gift from Didier Trono, Addgene plasmid #12260) plasmids into the HEK293T packaging cell line, using polyfectamine reagent (Qiagen), and incubated at 37°C to generate lentiviral particles. After 48 h, the culture supernatant was passed through a 0.45 μm filter to harvest viral particles, and infect U-2 OS cells following treatment with polybrene (8 μg/mL, Sigma-Aldrich) for 2 h. 2 μg/mL puromycin (Sigma-Aldrich) was added to cells after 48 h to select transduced cells. The extent of knockdown was determined by immunoblot analysis with an IP6K1 specific antibody. For transfection, polyethylenimine (PEI) (Polysciences, 23966) was used at a ratio of 1:3 (DNA:PEI). All plasmids used for transfection were purified using the Plasmid Midi kit (Qiagen). Cells were harvested 36-48 h post-transfection for further analyses.

### Mice

All animal experiments were approved by the Institutional Animal Ethics Committee (Protocol number EAF/CDFD/RB/01), and were performed in compliance with guidelines provided by the Committee for the Purpose of Control and Supervision of Experiments on Animals, Government of India. *Ip6k1^+/+^* and *Ip6k1^−/−^* mice *(Mus musculus*, strain C57BL/6) used for this study were housed in the Experimental Animal Facility at the Centre for DNA Fingerprinting and Diagnostics, Hyderabad. *Ip6k1^+/+^* and *Ip6k1^−/−^* littermates were generated and maintained as previously described (5).

### Immunofluorescence analysis

Cells grown on glass coverslips placed in a 24 well plate were fixed with 4% paraformaldehyde (PFA) for 10 min at room temperature (RT), and permeabilized with 0.15% Triton-X 100 for 10 min at RT. Non-specific antibody binding was blocked by incubating the cells in a blocking buffer (3% BSA in PBS with 0.1% Triton-X 100) for 1 h at RT. After blocking, primary antibody diluted in the blocking buffer was added to the cells, and incubated overnight at 4°C. Cells were washed with PBS three times, incubated with fluorophore conjugated secondary antibodies (Alexa Fluor 488 or Alexa Fluor 568 goat anti-rabbit or anti-mouse IgG; 1:500) diluted in the blocking buffer, and incubated for 1 h at RT. Cells were washed with PBS three times, and the coverslips were mounted on glass slides using an antifade mounting medium with DAPI (H-1200, Vecta Labs), air dried and sealed. For P-body staining of bronchiolar epithelium tissue, paraffin embedded sections from *Ip6k1*^+/+^ and *Ip6k1*^−/−^ mice were treated with xylene for deparaffinization, dipped in a series of graded ethanol for dehydration, and boiled in 10 mM sodium citrate buffer, pH 6, for 10 min to achieve antigen retrieval. The sections were then washed, permeabilized, and stained as detailed above. Images were acquired on a Zeiss LSM 700 confocal microscope equipped with 405, 488 and 555 nm lasers, and fitted with a 63x, 1.4 NA objective. The number of P-bodies per cell was quantified using Fiji software (37), as described earlier (38). Briefly, Z-stacks (step size 0.5 μm) were digitally collapsed, and the threshold for P-body quantification was set using Otsu thresholding plugin. P-bodies were counted using the ‘analyse particles’ feature of Fiji, where pixel range to be counted as P-bodies was set at 7 - 500.

### Co-immunoprecipitation and western blot analysis

Transfected HEK293T cells were collected after 48 h, and lysed for 1 h at 4°C in cell lysis buffer (50 mM HEPES pH 7.4, 100 mM NaCl, 1 mM EDTA, 0.5% Nonidet P40, protease inhibitor cocktail (Sigma P8340), and phosphatase inhibitor cocktail (Sigma P5762)). Specific antibody was added to the lysate and incubated overnight at 4°C with rotation, and the complex was pulled down using pre-equilibrated Protein A Sepharose beads (GE Healthcare) for 1 h. The beads were washed three times with lysis buffer, boiled in 1X Laemmli buffer, and processed using standard western blotting techniques. Chemiluminescence was detected using the UVITEC Alliance Q9 documentation system, or the GE ImageQuant LAS 500 imager. Densitometry analysis of bands was done using Fiji software (37). In case of HEK293T cells expressing SFB tagged proteins, pre-equilibrated streptavidin sepharose beads were added to the cell lysate, and incubated for 90 min at 4°C prior to washing and boiling the beads. In case of RNase experiments, RNase A was added to the lysate at 100 μg/mL for 1 h at 4°C before the addition of beads.

For m^7^GTP pull down experiments, cells were lysed in cap lysis buffer (50mM HEPES pH 7.4, 100mM NaCl, 1mM EDTA, 1% Nonidet P40, protease inhibitor cocktail (Sigma P8340), and phosphatase inhibitor cocktail (Sigma P5762)), and incubated with 20 μL of pre-equilibrated γ-aminophenyl-m^7^GTP beads (Jena Bioscience) at 4°C overnight. The beads were washed three times with cap lysis buffer, and proteins were detected by immunoblotting as described above. Where indicated, the lysates were incubated with 100 μM m^7^GpppG cap analogue at 4°C for 30 min prior to the addition of γ-aminophenyl-m^7^GTP beads.

### RT-qPCR

Total cellular RNA was isolated from U-2 OS NT or sh*IP6K1* cells using TRIzol reagent (Invitrogen) for cell lysis, and RNeasy Mini Kit (Qiagen) for RNA extraction. 2 μg RNA was used to synthesize cDNA by reverse transcription with oligonucleotide dT primers using SuperScript Reverse Transcriptase III (Invitrogen). Gene-specific primers were used to perform qPCR on the ABI 7500 Real-Time PCR System (Applied Biosystems) with MESA GREEN qPCR MasterMix Plus and SYBR® Assay Low ROX (Eurogentec) for detection. Each sample was run in technical duplicates. The sequences of primers used are listed in supplemental table S1. The difference in transcript levels was calculated using the fold change (ΔΔC_t_) method (39). ΔC_t_ is the C_t_ value for the gene of interest normalized to the C_t_ value of the respective GAPDH control in both U-2 OS NT and U-2 OS sh*IP6K1* cells. ΔΔC_t_ values were calculated as a relative change in ΔC_t_ of the target gene in U-2 OS sh*IP6K1* cells with respect to U-2 OS NT cells. Fold changes were expressed as 2^− ΔΔCt^.

### Subcellular fractionation of U-2 OS and HEK293T cells

~2 ×10^7^ cells were trypsinized, washed with 1X PBS, lysed by resuspension in Buffer A (250 mM sucrose, 250 mM KCl, 5 mM MgCl_2_, 50 mM Tris·Cl, pH 7.4, and 0.7% Nonidet-P40), and centrifuged at 750 *g* to pellet the nuclear fraction. The post-nuclear supernatant was further centrifuged at 12,500 *g* to pellet the mitochondrial fraction. The concentration of KCl in the post-mitochondrial fraction (PMF) was adjusted to 0.5 M to disrupt weak interactions between proteins of other cellular compartments and the ribosomes. Finally, to obtain a ribosomal pellet, the PMF was layered on top of a sucrose cushion (1 M sucrose, 0.5 M KCl, 5 mM MgCl_2_, and 50 mM Tris Cl, pH 7.4), and subjected to ultracentrifugation at 250,000 *g* at 4°C for 2 h. The ribosomes were present in a translucent and viscous pellet. The post-nuclear, ribosomal, and post-ribosomal fractions were boiled in 1X Laemmli buffer, and processed using standard western blotting techniques. For immunoprecipitation from ribosomes, the ribosomal pellet was resuspended in cell lysis buffer, and processed as described above.

### Luciferase based miRNA reporter assay

The luciferase reporters used in this assay are described in (35), and were obtained from Dr. Suvendra Bhattacharyya (CSIR-Indian Institute of Chemical Biology, Kolkata). HeLa NT and HeLa sh*IP6K1* cells were transfected in a 24-well plate with 100 ng of pFL (firefly luciferase) transfection control plasmid, and 75 ng of pRL (Renilla luciferase) constructs (pRL control, pRL-perf, or pRL-3XB). 24 h post transfection, luciferase assay was performed using the Dual Luciferase Reporter Assay kit (Promega) as per the manufacturer’s instructions. Luciferase activity was measured using the EnSpire Multimode Plate Reader (PerkinElmer). Renilla luciferase activity was normalized to firefly luciferase activity for each sample. These normalized values were expressed relative to the pRL control sample for each cell line.

### Decapping assay

The mRNA decapping assay was performed as previously described (40). Briefly, a 176-bp luciferase cDNA fragment, amplified from pGL3-control plasmid (Promega), was transcribed and capped using the T7 mScript™ Standard mRNA Production System (CellScript). The RNA was labelled by adding 20 μCi of [α-^32^P]GTP to the capping reaction. The RNA was purified after each step - transcription and capping – using the total RNA isolation protocol outlined with the mirVana miRNA isolation kit (Invitrogen). To obtain GFP - tagged DCP2 for the decapping assay, HEK293T cells transfected with plasmids encoding GFP-DCP2, or the GFP control, with or without IP6K1-V5, were lysed in NET-1 buffer (50 mM Tris pH 7.5, 150 mM NaCl, 1 mM EDTA, 0.1% Triton X-100, and 10% glycerol). GFP fusion proteins were immunoprecipitated as described above, with the exception that the last washing step was performed with NET-2 buffer (50 mM Tris HCl, pH 7.5, 150 mM NaCl, 0.05% Triton X-100, and 0.1 mg/mL BSA, supplemented with protease and phosphatase inhibitor cocktail). Immunoprecipitates on Protein A Sepharose beads were resuspended in NET-2 buffer to form a 1:1 slurry. Half the immunoprecipitate was used for immunoblotting, and the remaining half was used for the decapping assay. The immunoprecipate was resuspended in decapping buffer (50 mM Tris-HCl, pH 7.9, 30 mM ammonium sulfate, and 1 mM MgCl_2_), containing 0.1 mM cap structure analogue m^7^GpppG (Jena Bioscience), 0.4 U/μL RNase inhibitor, and 10^5^ cpm of capped RNA, in a 15 μL reaction volume, and incubated for 30 min at 30°C with mixing at 900 rpm (Thermomixer R, Eppendorf). The reaction was stopped by the addition of 50 mM EDTA, 7 μL sample was spotted on a polyethylenimine (PEI) cellulose TLC plate (Sigma-Aldrich, Z122882), and resolved using 0.75 M LiCl. Unlabeled m^7^GMP, m^7^GDP, and m^7^GTP were used as standards, and were visualized by UV shadowing. The radiolabelled capped mRNA substrate and m^7^GDP product were visualised by autoradiography (Typhoon FLA-9500, GE). The extent of decapping was measured by normalizing the intensity of the m^7^GDP product to the intensity of the capped mRNA substrate spotted at the origin.

### Statistical analysis

GraphPad Prism 5 was used to perform statistical analyses and prepare graphs. Densitometry data for western blots were obtained using Fiji software (37). Band intensities of indicated proteins were normalized to their respective loading controls from the same blot. These normalized intensity values were expressed relative to the control in each blot, as detailed in the figure legends. The number of cells (n) used to perform statistical tests for each experiment, and the number of biologically independent replicates (N) for each experiment are indicated in the figure legends. *P* values are from either a one-sample *t*-test or a two-tailed unpaired Student’s *t*-test, as indicated in the respective figure legends. *P* ≤ 0.05 was considered statistically significant.

## Acknowledgements

The authors thank Suvendra Bhattacharyya for generously sharing Renilla luciferase miRNA reporters. We thank Aushaq Bashir Malla for initiating investigations into the effect of IP6K1 on P-bodies; Shubhra Ganguli for the generation of U-2 OS NT and sh*IP6K1* cells and for the construction of SFB-IP6K1 expression plasmid; Vineesha Oddi for the construction of IP6K1-V5 expression plasmid; and Aisha Hamid for the construction of IP6K1-V5 K226A expression plasmid. We acknowledge the staff at the Sophisticated Equipment Facility and Experimental Animal Facility, CDFD, for technical assistance. We thank Maddika Subba Reddy, Rohit Joshi, and members of the Laboratory of Cell Signalling for valuable feedback.

## Author contributions

Conceived research: A.S. and R.B.; Designed and performed the experiments: A.S.; Data analysis: A.S. and R.B.; Wrote the paper: A.S. and R.B.

