## Supplementary Figures and Table for "IP6K1 upregulates the formation of processing bodies by promoting proteome remodeling on the mRNA cap"

**This PDF file includes:**

Supplementary Figures S1 to S3
Supplementary Table S1

#### Figure S1

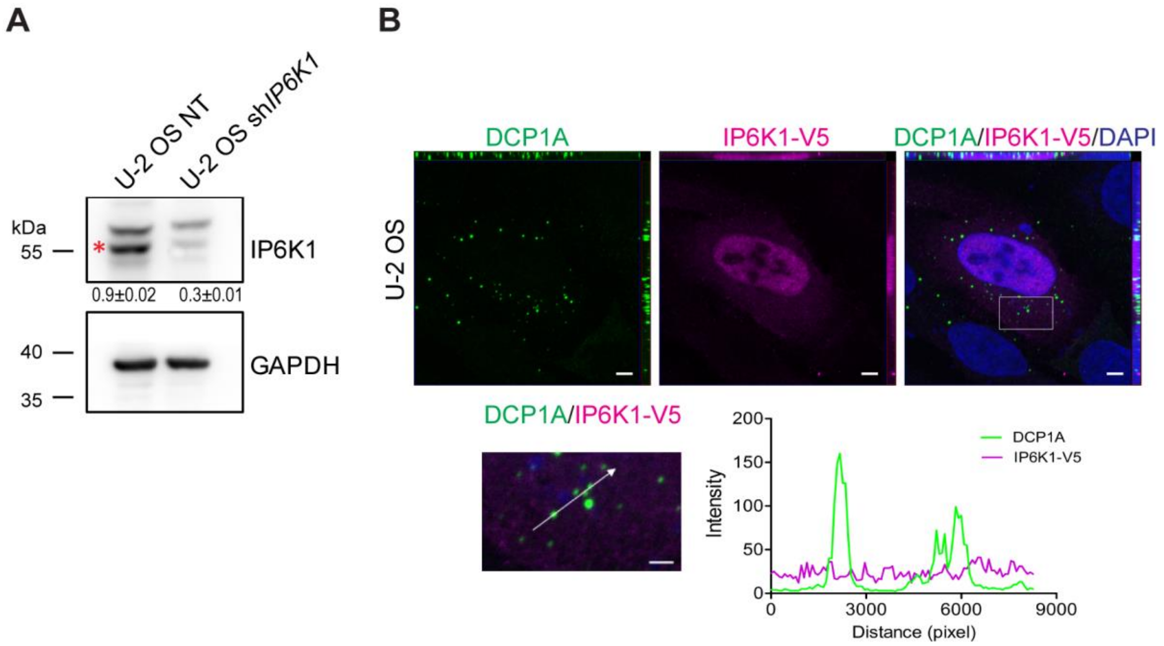


**Figure S1. Overexpressed IP6K1 does not localize to P-bodies. (A)** Representative immunoblot to detect IP6K1 in U-2 OS cells stably expressing either non-targeting control (NT), or shRNA directed against human IP6K1 (sh*IP6K1*). Numbers show mean fold change ± SEM, N=4. This data is also shown as a bar graph in Fig. 3B. **(B)** Localization of V5 epitope-tagged IP6K1 (magenta) with DCP1A (green) in U-2 OS cells. Nuclei were stained with DAPI (blue). Scale bar, 5 μm, N=3. The boxed region in (B) is magnified and shown below, where intensity profiles were measured along the line drawn. Scale bar, 2 μm. The green and magenta traces denote DCP1A and IP6K1 fluorescence intensities, respectively, and the high intensity green peaks correspond to P-bodies.

### Figure S2

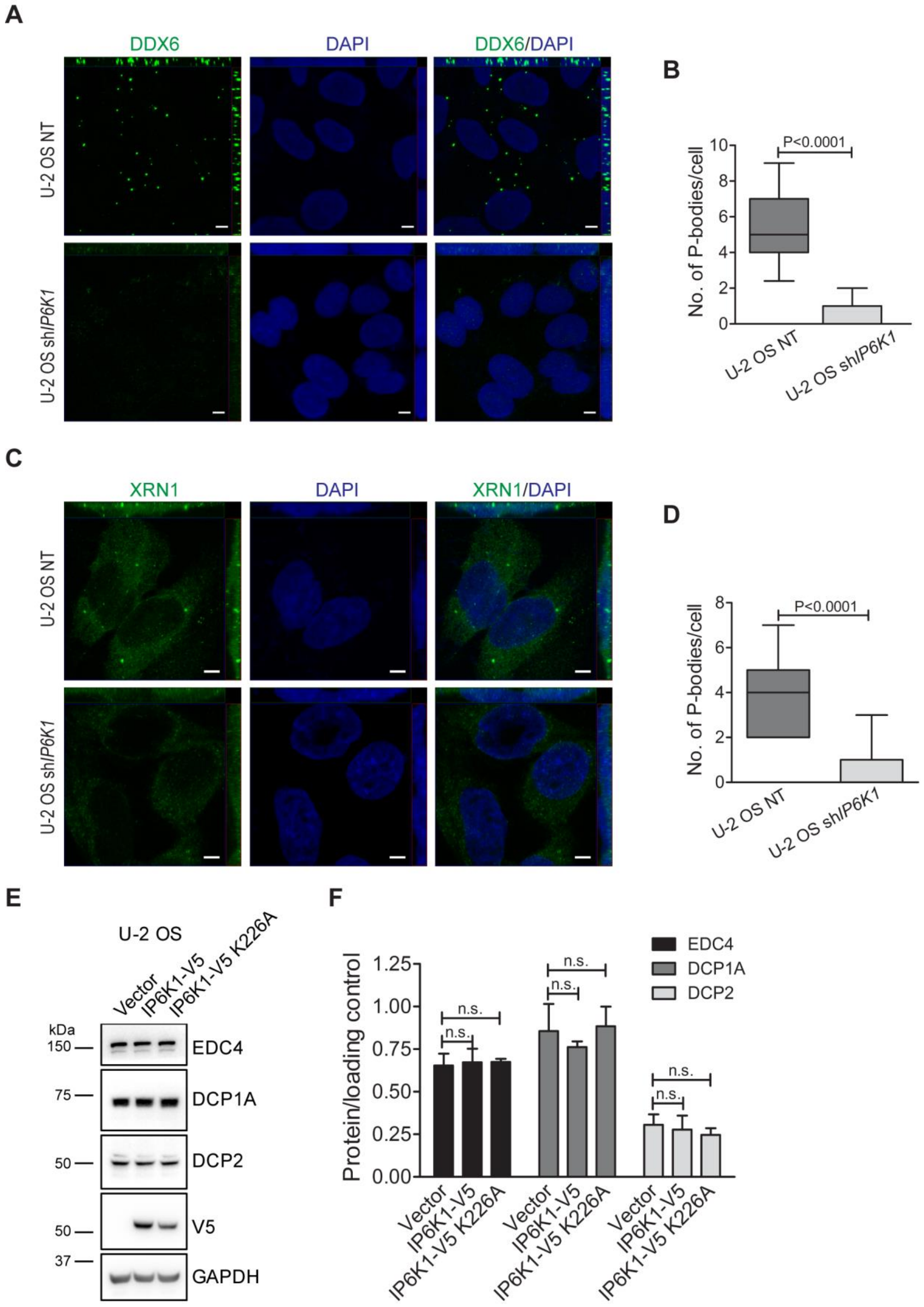
A
DDX6
DAPI
DDX6/DAPI
U-2 OS NT
U-2 OS shIP6K1
B
No. of P-bodies/cell
P<0.0001
U-2 OS NT
U-2 OS shIP6K1
C
XRN1
DAPI
XRN1/DAPI
U-2 OS NT
U-2 OS shIP6K1
D
No. of P-bodies/cell
P<0.0001
U-2 OS NT
U-2 OS shIP6K1
E
U-2 OS
Vector
IP6K1-V5
IP6K1-V5 K226A
kDa
150
75
50
50
37
EDC4
DCP1A
DCP2
V5
GAPDH
F
Protein/loading control
n.s.
EDC4
DCP1A
DCP2
Vector
IP6K1-V5
IP6K1-V5 K226A

**Figure S2. DDX6 and XRN1 stained P-bodies are depleted from cells with reduced IP6K1 expression. (A)** Asynchronous U-2 OS NT and sh*IP6K1* cells were stained with anti-DDX6 antibody (green). Nuclei were stained with DAPI (blue). Scale bar, 5 µm. **(B)** Quantification of the number of P-bodies/cell in (A). Data (mean ± SEM, n = 91 and 119 cells respectively, for U-2 OS NT and sh*IP6K1* cells) are representative of two independent experiments. **(C)** Asynchronous U-2 OS NT and sh*IP6K1* cells were stained with anti-XRN1 antibody (green). Nuclei were stained with DAPI (blue). Scale bar, 5 µm. **(D)** Quantification of the number of P-bodies/cell in (C). Data (mean ± SEM, n = 51 and 71 cells respectively, for U-2 OS NT and sh*IP6K1* cells) are representative of two independent experiments. **(E)** Representative immunoblots showing the levels of mRNA decapping proteins in U-2 OS cells overexpressing either IP6K1 or IP6K1 K226A. GAPDH was used as a loading control. **(F)** Protein levels from (E) were normalized to their respective loading control. Data are mean ± SEM, N=3. Images in (A and C) were subjected to uniform 'levels' adjustment in the ZEN software to improve visualization. *P* values are from a two-tailed unpaired Student's *t*-test (B, D and F); $P \leq 0.05$ was considered significant, n.s., not significant $P > 0.05$.

Figure S3

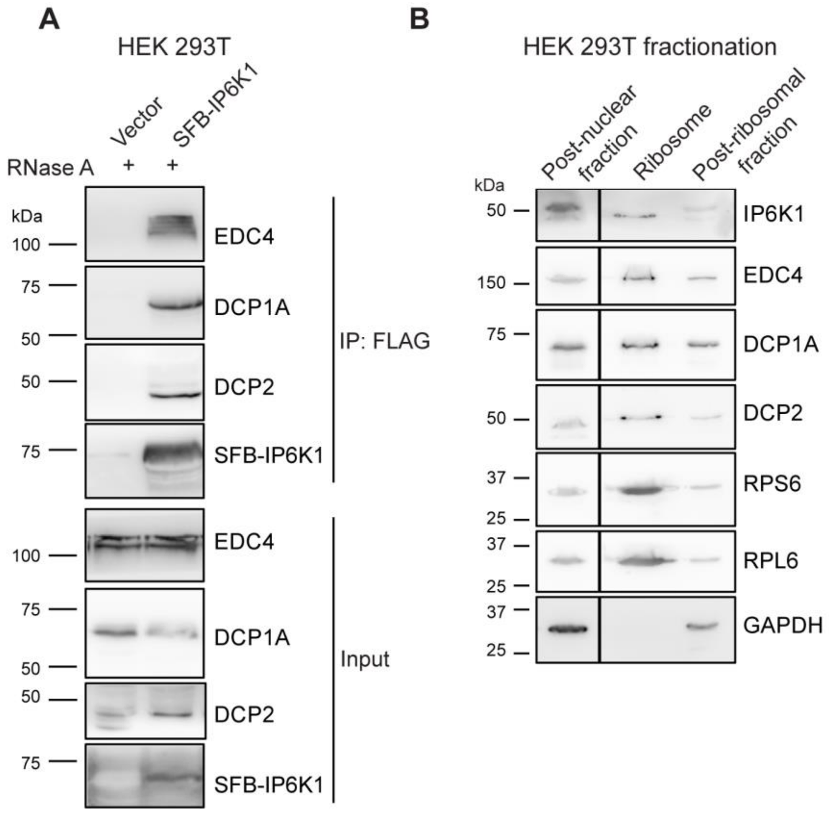


**Figure S3. Interaction of IP6K1 with mRNA decapping proteins is RNA independent. (A)** Representative immunoblots showing co-immunoprecipitation of endogenous DCP1A, EDC4, and DCP2 with SFB-tagged IP6K1 from RNase A-treated cell extracts. SFB-IP6K1 was transiently overexpressed in HEK293T cells, immunoprecipitated with an anti-FLAG antibody, and probed to detect DCP1A, EDC4, or DCP2. The SFB tag was detected using an anti-FLAG antibody (N=3). **(B)** Representative immunoblots of subcellular fractions of HEK293T cells prepared by ultracentrifugation, to detect endogenous IP6K1, EDC4, DCP1A, and DCP2. Enrichment of ribosomes was marked by the presence of RPL6 and RPS6, and GAPDH was detected to rule out cytoplasmic contamination in the purified ribosomes (N=2). Vertical line indicates removal of non-essential lanes from a single original gel to improve visualisation.

**Table S1:** Primers used for RT-qPCR analysis shown in Fig. 3C

| Transcript | Primer |
|---|---|
| *DCP1A* | F: 5’- AATAAAGAATGATTCCAGCTTCCTC -3’<br>R: 5’- AGCCTATTTGTCTCTGAGGCTG -3’ |
| *DCP1B* | F: 5’- TATGTCTGGGAGGGAGGGAAG -3’<br>R: 5’- CACATCAGTTTTCTCCCACTCGT -3’ |
| *DCP2* | F: 5’- TGGAAGGTTGTTCAGATACTGGC -3’<br>R: 5’- AGAGCAACCACAATGAGCAAC -3’ |
| *EDC4* | F: 5’- ATTACAAGGGCCGATGCAGG -3’<br>R: 5’- GCTGCTGCAAGTATTCCTGTG -3’ |
